# HOIL1 mediates MDA5 activation through ubiquitination of LGP2

**DOI:** 10.1101/2024.04.02.587772

**Authors:** Deion Cheng, Junji Zhu, GuanQun Liu, Michaela U. Gack, Donna A. MacDuff

## Abstract

The RIG-I-like receptors (RLRs), RIG-I and MDA5, are innate sensors of RNA virus infections that are critical for mounting a robust antiviral immune response. We have shown previously that HOIL1, a component of the Linear Ubiquitin Chain Assembly Complex (LUBAC), is essential for interferon (IFN) induction in response to viruses sensed by MDA5, but not for viruses sensed by RIG-I. LUBAC contains two unusual E3 ubiquitin ligases, HOIL1 and HOIP. HOIP generates methionine-1-linked polyubiquitin chains, whereas HOIL1 has recently been shown to conjugate ubiquitin onto serine and threonine residues. Here, we examined the differential requirement for HOIL1 and HOIP E3 ligase activities in RLR-mediated IFN induction. We determined that HOIL1 E3 ligase activity was critical for MDA5-dependent IFN induction, while HOIP E3 ligase activity played only a modest role in promoting IFN induction. HOIL1 E3 ligase promoted MDA5 oligomerization, its translocation to mitochondrial-associated membranes, and the formation of MAVS aggregates. We identified that HOIL1 can interact with and facilitate the ubiquitination of LGP2, a positive regulator of MDA5 oligomerization. In summary, our work identifies LGP2 ubiquitination by HOIL1 in facilitating the activation of MDA5 and the induction of a robust IFN response.

## Introduction

The RIG-I-like receptors (RLRs), RIG-I and MDA5, are major innate sensors of RNA virus infections and initiate antiviral immunity through the induction of type 1 interferons (IFN) and inflammatory cytokines (1, 2). RIG-I is activated by short dsRNAs with 5’ tri- or di-phosphate and is the major sensor for many negative-sense RNA viruses, including orthomyxoviruses, paramyxoviruses, and rhabdoviruses (3, 4). MDA5 is primarily activated by long dsRNAs and is critical for innate sensing of coronaviruses, picornaviruses and caliciviruses (4–7). Upon binding their respective RNA ligands, RIG-I and MDA5 oligomerize and signal to the outer mitochondrial membrane protein, MAVS, via an interaction between their respective caspase activation and recruitment domains (CARDs). This interaction triggers MAVS to aggregate into a signaling platform that recruits IRF3 or IRF7 and NF-κ;B transcription factors and their activating kinases (TBK1, IKKε, IKKα and IKKβ), leading to the upregulation of IFN, IFN-stimulated genes (ISGs) and pro-inflammatory cytokines (2, 8).

LGP2 is the third member of the RLR family. LGP2 has RNA-binding capability but lacks CARDs to initiate signaling to MAVS. Instead, the role of LGP2 in antiviral signaling is to modulate the activities of RIG-I and MDA5 (9–13). LGP2 has been shown to both positively and negatively regulate RLR signaling, with these opposing roles appearing to depend on LGP2 expression levels (9). Nevertheless, LGP2 deficiency results in a near complete loss of an IFN response upon infection with viruses that require MDA5 sensing, indicating an essential role for LGP2 in MDA5 antiviral signaling (12, 13). At the molecular level, LGP2 was shown to facilitate MDA5 oligomer formation along dsRNA (9, 14).

Beyond regulation by LGP2, MDA5 and RIG-I activation and signaling to MAVS are each controlled by distinct networks of protein interactions, post-translational modifications, and other regulatory mechanisms (2). Such complex regulation allows for tight control of RLR-driven immune responses during infection to safeguard against potentially detrimental interferonopathies and auto-inflammatory disorders caused by overactive signaling (15–19). Therefore, a thorough understanding of the molecular events governing RLR activation and signaling holds promise for the development of host-directed antivirals or targeted anti-inflammatory therapies.

We previously showed that Heme-oxygenase IRP2 ligase 1 (HOIL1, also known as RBCK1) is required for MDA5 antiviral signaling in cells and to control norovirus infection in mice, but has no apparent role in RIG-I signaling (20). However, the functional contribution of HOIL1 remained ambiguous due to its integral role as a part of the linear ubiquitin chain assembly complex (LUBAC). The LUBAC is a trimeric complex consisting of HOIL1, HOIL1-Interacting Protein (HOIP, also known as RNF31), and SHANK-associated RH domain interactor (SHARPIN), that generates linear (methionine-1-linked) polyubiquitin chains (21–24). Linear ubiquitination regulates multiple immune-related pathways, such as canonical NF-κB activation, NLRP3 inflammasome formation, and extrinsic cell death pathways (21, 22, 25–32). Several studies have also shown that the LUBAC negatively RIG-I signaling, while other studies have reported an opposing role (33–37).

HOIP functions as the E3 ubiquitin ligase that catalyzes linear ubiquitination, while HOIL1 and SHARPIN promote LUBAC stability and allosterically activate HOIP (21–25, 38, 39). Besides supporting LUBAC stability, HOIL1 is also a functional E3 ubiquitin ligase (39–42). HOIL1 ligase activity is comparatively understudied, though emerging evidence has demonstrated that HOIL1 catalyzes atypical ester-linked ubiquitination of serine and threonine residues as well as glucose of unbranched polysaccharides (41–47). Recent studies also found coordinated activity between HOIL1 and HOIP ligase activities in the form of heterotypic ubiquitin chains conjugated onto the same substrate protein, although the biological implications for such modifications are largely unknown. Thus, whether HOIL1 or HOIP E3 ligase activity, or both, regulate MDA5 signaling is undetermined.

Here, we examined the relative contributions of HOIL1 versus HOIP E3 ligase activity to RLR signaling and IFN induction during RNA virus infection. We demonstrate that HOIL1 E3 ubiquitin ligase activity is selectively necessary for MDA5 antiviral signaling and facilitates the oligomerization of MDA5, leading to MAVS activation. In contrast, HOIP catalytic activity promotes both MDA5 and RIG-I signaling in a distinct and more minor capacity. Finally, we investigated the functional contribution of HOIL1 ligase activity to MDA5 signaling and identified LGP2 as a link between HOIL1 and MDA5.

## Results

### HOIL1 E3 ligase activity is critical for MDA5 antiviral signaling

We have shown previously that HOIL1 is essential for IFN induction after infection with viruses that activate MDA5 (20). However, since HOIL1 deficiency significantly impairs the stability and function of its binding partners, HOIP and SHARPIN, we were unable to establish whether HOIL1 facilitates MDA5-dependent IFN induction through its intrinsic E3 ubiquitin ligase activity, or through regulation of HOIP and SHAPRIN. To examine the requirement for HOIL1 catalytic activity, we stably complemented wild-type (WT) and HOIL1-deficient (*Hoil1*^-/-^) mouse embryonic fibroblasts (MEFs; from B6NCrl;B6N-A^tm1Brd^Rbck1^tm1a(EUCOMM)Hmgu^ mice (48)) with FLAG-tagged wildtype HOIL1 (HOIL1[WT]), HOIL1 bearing an inactivating point mutation in the ubiquitin ligase domain (C458S or C458A), or a firefly luciferase (FLUC) negative control. In line with previous studies (41–43), HOIL1[C458S] and HOIL1[C458A] were able to complex with LUBAC components HOIP and SHARPIN, similar to HOIL1[WT] (**Fig. 1A**). Furthermore, while HOIL1 deficiency sensitized the MEFs to TNFα-induced cell death due to impairment of HOIP expression and activity (49), HOIL1[C458S] and HOIL1[C458A] restored HOIP and SHARPIN expression and resistance to TNFα-induced cell death, similar to HOIL1[WT], indicating that ligase-dead HOIL1 can support HOIP catalytic activity (**Fig. 1A**).

**Figure 1:**
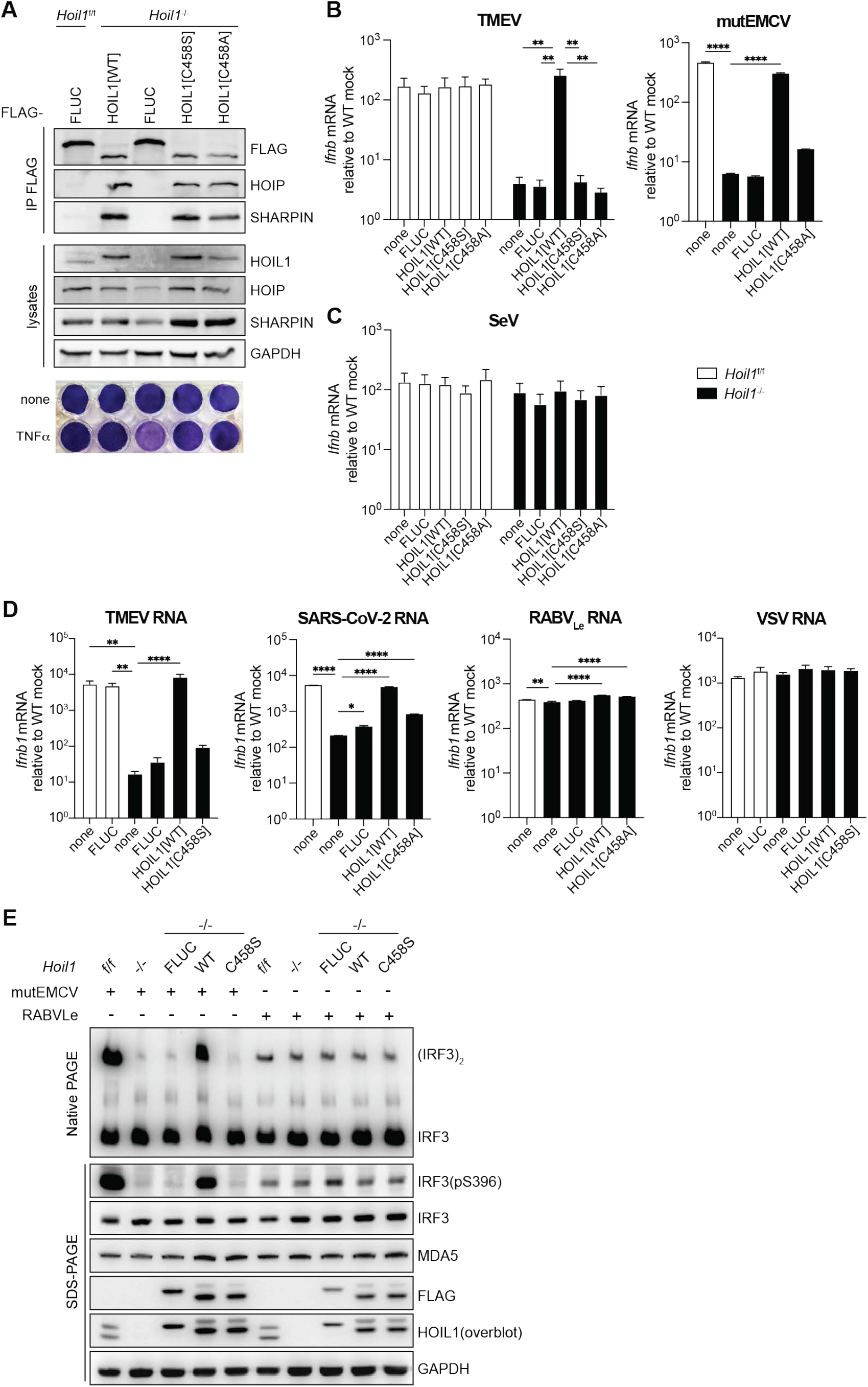
HOIL1 E3 ubiquitin ligase activity is essential for MDA5 signaling and *Ifnb1* induction. *Hoil1*^f/f^ (WT) and *Hoil1*^-/-^ MEFs were stably transduced with vectors expressing 3xFLAG-HOIL1[WT], -HOIL1[C458S], -HOIL1[C458A], –FLUC, or untransduced (none). (**A**) Immunoblot analyses of anti-FLAG co-IPs and of whole cell lysates (top and middle), and crystal violet stain of cells treated with 150 µg/ml TNFα for 48 h (bottom), from the indicated MEF lines. (**B-C**) *Ifnb1* mRNA fold induction in the indicated MEFs in response to infection with TMEV (MOI of 10, 11 hpi), mutEMCV (MOI of 0.5, 12 hpi), SeV (MOI of 3, 9 hpi), expressed as relative to mock-infected WT cells. (**D**) *Ifnb1* mRNA fold induction in the indicated MEFs after transfection with the indicated viral RNAs: TMEV RNA (500 μg/ml) for 8 h; VSV RNA (500 μg/ml) for 8 h, SARS-CoV-2 RNA (400 ng/ml) for 16 h or RABV_Le_ RNA (2 pmol/ml) for 16 h. (**E**). Native PAGE analysis of IRF3 dimerization (top) and SDS-PAGE analysis of IRF3 phosphorylation, and expression of the indicated proteins in MEFs following infection with mutEMCV (MOI of 5) or transfection with RABV_Le_ RNA (5 pmol/ml) for 8 h. For A and E, data shown are representative of at least 3 independent experiments. For B-D, TMEV, SeV, TMEV RNA and VSV RNA data are combined means from 3-4 independent experiments performed in duplicate. mutEMCV, SARS-CoV-2 RNA and RABV_Le_ RNA data are representative of at least two independent experiments, and the mean ± SEM is shown. Significance was determined by two-way ANOVA with Tukey’s multiple comparisons test (B, C), or by one-way ANOVA compared to *Hoil1*^-/-^ untransduced with Dunnett’s multiple comparisons test (D). \**p*≤0.05, \*\**p*≤0.01, \*\*\**p*≤0.001, \*\*\*\**p*≤0.0001.

To determine whether HOIL1 ligase activity regulates IFN induction after MDA5 activation, the aforementioned complemented MEFs were infected with viruses that predominantly activate MDA5 [i.e., Theiler’s murine encephalomyelitis virus (TMEV) and encephalomyocarditis virus (mutEMCV, a mutant strain deficient in antagonism of MDA5 (50)] or RIG-I [Sendai virus (SeV)], and *Ifnb1* mRNA induction measured. As observed previously, *Hoil1*^-/-^ cells exhibited a 42-fold and 75-fold reduction in *Ifnb1* induction after infection with TMEV and mutEMCV, respectively (**Fig. 1B**) (20). While HOIL1[WT] restored MDA5-driven *Ifnb1* induction in *Hoil1*^-/-^ MEFs, HOIL1[C458S/A] failed to restore *Ifnb1* mRNA. In contrast, HOIL1 deficiency or expression of HOIL1[C458S/A] did not significantly impact *Ifnb1* induction upon RIG-I stimulation by SeV infection, as expected (**Fig. 1C**). Similar results were observed upon transfection of MEFs with immunostimulatory viral RNAs (51) that predominantly activate MDA5 (TMEV or SARS-CoV-2 RNA) or RIG-I (RABV leader [RABV_Le_] RNA or VSV RNA), further supporting the requirement for HOIL1’s E3 ligase activity in promoting MDA5, but not RIG-I, signaling (**Fig. 1D**). In accord, IRF3 phosphorylation and dimerization were severely impaired in HOIL1[C458S]-reconstituted MEFs following MDA5 activation by mutEMCV infection, but not following RIG-I activation by RABV_Le_, demonstrating that HOIL1 E3 ligase activity functions upstream of IRF3 activation in the MDA5 signaling pathway (**Fig. 1E**).

We further corroborated these findings by measuring the IFN response of bone-marrow derived dendritic cells (BMDCs) generated from *Hoil1*^C458S/C458S^ knock-in mice (41), with *Mda5*^-/-^ BMDCs as controls. In agreement with results obtained from transduced MEFs, BMDCs expressing HOIL[C458S] exhibited ∼100-fold reduced *Ifnb1* mRNA expression after infection with murine norovirus (MNoV) or TMEV, but not with SeV or VSV (**Fig. 2A**). In comparison, MDA5-deficient BMDCs exhibited a ∼1000-fold reduction in *Ifnb1* mRNA expression after MNoV or TMEV infection, indicating that HOIL1 E3 ligase activity is important, but not absolutely required, for MDA5-mediated IFN induction (**Fig. 2A**). Similar results were observed for the transcriptional induction of *Ifit2* (an IRF3-induced ISG) and *Il6* (a major proinflammatory cytokine), indicating that activation of both IRF3 and NF-κB is impacted by loss of HOIL1 ligase activity (**Fig. 2B, C**). However, BMDCs from *Hoil1*^C458S/C458S^ mice did not exhibit defects in *Ifnb1* or *Il6* induction in response to LPS treatment, which activates IRF3 and NF-κB through activation of TLR4 (**Fig. 2D**). Instead, *Ifnb1* mRNA was elevated in cells expressing HOIL1[C458S], which is consistent with reports indicating that HOIL1 E3 ligase inhibits linear ubiquitination (41, 42, 52). Importantly, HOIP and SHARPIN were expressed at similar levels in WT and *Hoil1*^C458S/C458S^ cells (**Fig. 2E**). Thus, unlike complete HOIL1 deficiency, targeted mutation of HOIL1 ligase allowed us to disentangle the role of HOIL1 E3 catalytic activity from stabilization of the LUBAC and HOIP protein levels. Together, our data establish HOIL1 E3 ubiquitin ligase activity as a critical positive regulator of MDA5 antiviral signaling.

**Figure 2:**
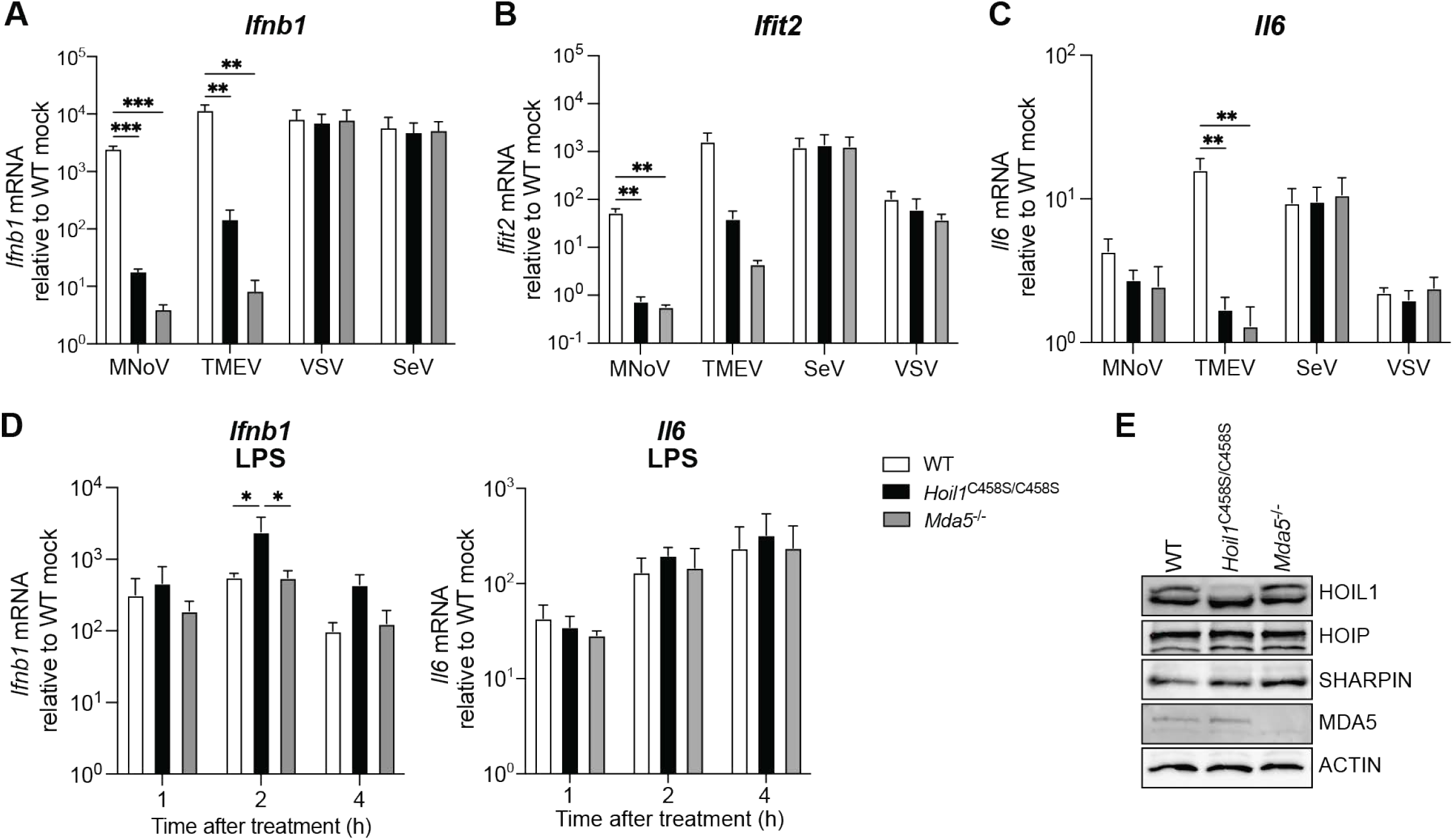
HOIL1 ligase activity is required in BMDCs for IRF3- and NF-κB-regulated gene induction in response to MDA5 agonists, but not RIG-I or TLR4 agonists. (**A-C**) Fold induction of *Ifnb1* (A), *Ifit2* (B) and *Il6* (C) mRNA in BMDCs from WT, *Hoil1*^C458S/C458S^, and *Mda5*^-/-^ mice infected by TMEV (MOI of 3, 9 hpi), MNoV (MOI of 3, 9 hpi), SeV (MOI of 0.3, 9 hpi), VSV (MOI of 3, 8 hpi), relative to WT mock-infected cells. (**D**) Fold induction of *Ifnb1* and *Il6* mRNA in BMDCs from WT, *Hoil1*^C458S/C458S^, and *Mda5*^-/-^ mice at the indicated times post-treatment with LPS. (**E**) Immunoblot analysis of the indicated proteins in uninfected BMDCs from WT, *Hoil1*^C458S/C458S^, and *Mda5*^-/-^ mice. For A-C and D, data are combined means from 3-4 independent experiments performed in duplicate, and the mean ± SEM is shown. Significance was determined by one-way ANOVA with Tukey’s multiple comparisons test (A-C), or by two-way ANOVA with Tukey’s multiple comparisons test (D). \**p*≤0.05, \*\**p*≤0.01, \*\*\**p*≤0.001.

### Differential requirements for HOIL1 and HOIP in RLR antiviral signaling

To investigate a possible interplay between HOIL1 and HOIP ligase activities in regulating MDA5 activation, we generated *Hoip*^-/-^ MEFs using CRISPR-Cas9-mediated targeting of exon 3 (encoding part of the N-terminal PUB domain) and confirmed gene disruption by sequencing. *Hoip*^-/-^ cells displayed greater sensitivity to TNFα-induced cell death than *Hoil1*^-/-^ cells, indicating severely impaired linear ubiquitination (**Fig. 3A**) (31, 53). Upon transfection with TMEV RNA, *Hoip*^-/-^ MEFs exhibited an approximately 5-fold defect in *Ifnb1* mRNA induction, compared to a 150-fold defect observed in *Hoil1*^-/-^ cells. To evaluate the requirement for HOIP E3 ligase activity, we stably expressed FLAG-tagged HOIP[WT] or its catalytically inactive mutants, HOIP[C879S] and HOIP[C879A] (38, 39, 54), in WT and *Hoip*^-/-^ cells. HOIP[C879S] and HOIP[C879A] were expressed to similar levels as HOIP[WT] in *Hoip*^-/-^ cells and formed complexes with HOIL1 and SHARPIN (**Fig. 3B**). While expression of HOIP[WT] restored resistance to TNFα-mediated cell death in *Hoip*^-/-^ cells, expression of HOIP[C879S] or HOIL1[C458A] did not, indicating successful inactivation of the HOIP catalytic site (**Fig. 3B**) (31).

**Figure 3:**
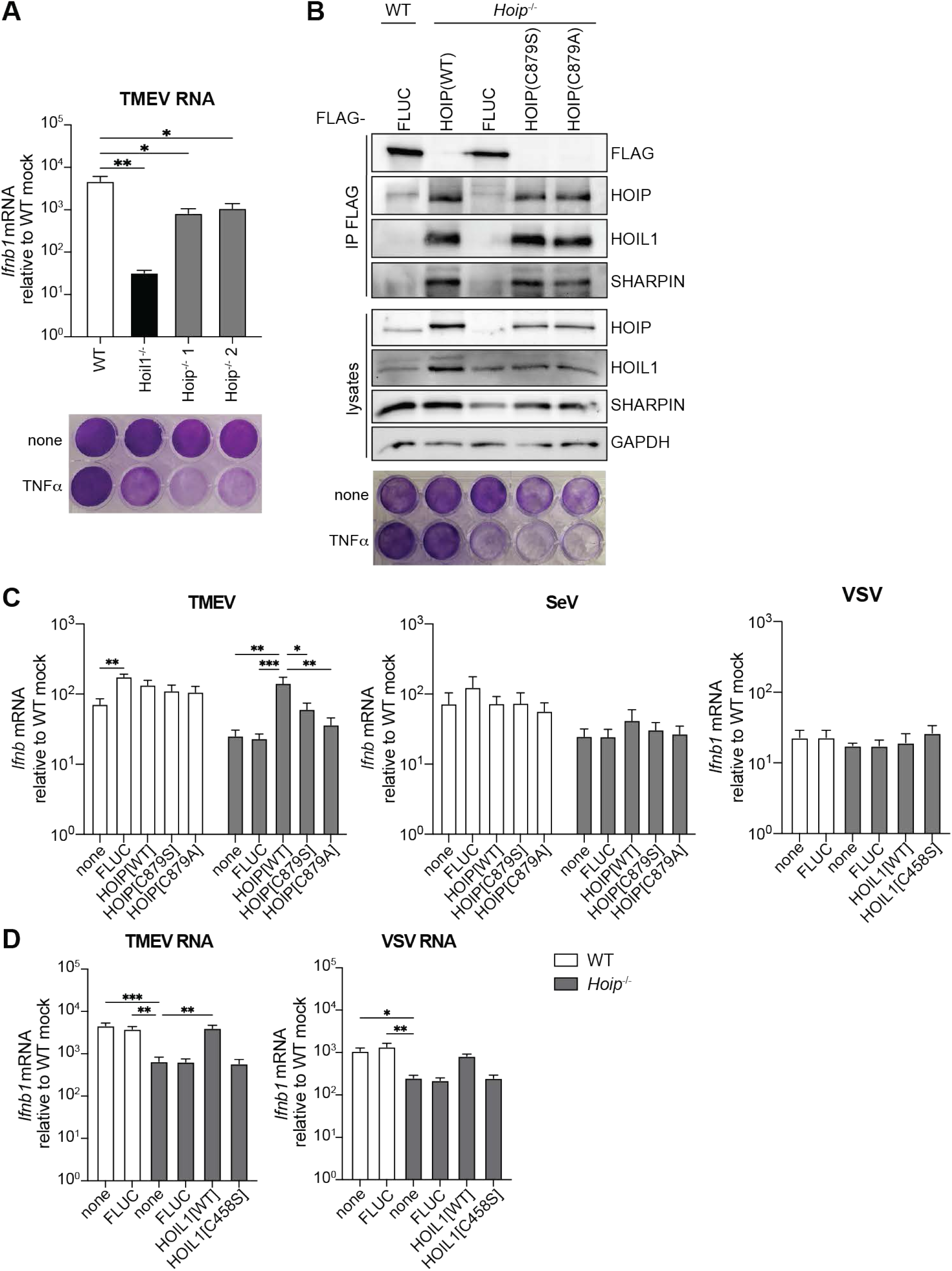
HOIP E3 ligase activity promotes both MDA5 and RIG-I-dependent *Ifnb1* induction. (**A**) *Ifnb1* mRNA induction after TMEV RNA transfection (top), and crystal violet-stained TNFα sensitivity assay (150 μg/ml for 48 h, bottom) for WT, *Hoil1*^-/-^ and *Hoip*^-/-^ MEFs. (**B-D**) WT and *Hoip*^-/-^ MEFs were stably transduced with vectors expressing 3xFLAG-HOIP[WT], -HOIP[C879S], -HOIP[C879A], –FLUC, or untransduced (none). (**B**) Immunoblot analyses of anti-FLAG co-IPs and whole cell lysates (top and middle), and crystal violet stain of cells treated with 150 μg/ml TNFα for 48 h (bottom), from the indicated MEF lines. (**C**) *Ifnb1* mRNA fold induction in the indicated MEFs after infection with TMEV (MOI of 10, 11 hpi), SeV (MOI of 3, 9 hpi), or VSV (MOI of 5, 8 hpi), relative to mock-infected WT cells. (**D**) *Ifnb1* mRNA fold induction in the indicated MEFs after transfection with the TMEV or VSV RNAs (500 μg/ml) for 8 h. For A (top), C and D, data are combined from 3-4 independent experiments performed in duplicate, and the mean ± SEM is shown. For A (bottom) and B, data shown are representative of 3 independent experiments. Significance was determined by one-way ANOVA with Tukey’s multiple comparisons test (A), two-way ANOVA with Tukey’s multiple comparisons test (C), or by one-way ANOVA compared to *Hoip*^-/-^ untransduced with Dunnett’s multiple comparisons test (D) and for VSV in (C). \**p*≤0.05, \*\**p*≤0.01, \*\*\**p*≤0.001.

To determine the role of HOIP catalytic activity in RLR signaling, the complemented *Hoip*^-/-^ MEFs were infected with TMEV, SeV or VSV (**Fig. 3C**). HOIP[WT] restored *Ifnb1* induction in *Hoip*^-/-^ cells after TMEV infection, but HOIP[C879S/A] did not. Although *Ifnb1* mRNA was slightly lower in *Hoip*^-/-^ cells after SeV infection, this was not significantly restored by HOIP[WT], and no defects in *Ifnb1* induction were observed after VSV infection. Since *Ifnb1* induction was generally low in these MEFs making it difficult to discern any differences, we transfected the MEFs with TMEV RNAs or VSV RNAs, which resulted in more robust IFN induction in WT cells. *Ifnb1* mRNA induction was reduced by 5-10-fold in *Hoip*^-/-^ cells transfected with TMEV RNAs (**Fig. 3D**), as before (**Fig. 3A**). While HOIP[WT] rescued *Ifnb1* induction, FLUC or HOIP[C458S] did not. However, comparable results were also observed upon transfection with VSV RNA (**Fig. 3D**), suggesting a role for HOIP E3 ligase activity in regulating both MDA5 and RIG-I pathways. The reduced IFN induction in response to VSV RNA may not have been observed in *Hoil1*^-/-^ MEFs due to partial stabilization of HOIP and SHARPIN likely by an N-terminal fragment of HOIL1 containing the UBL domain in those cells (**Fig. 1A, D**) (48). Together, these data indicate that HOIP E3 ligase activity plays a minor role in promoting RLR signaling and IFN induction, that may be obscured during viral infection, suggesting possible viral antagonism of RLR signaling or linear ubiquitination, or additional role(s) for HOIP in viral infection beyond innate RNA-sensing. These results corroborate findings from a recent study that reported a partial requirement for HOIP protein and HOIP E3 ligase activity for RIG-I signaling in A549 cells, reportedly through an interaction with and regulation of NEMO and TBK1 (37).

### HOIL1 regulates MDA5 signaling independent of its association with HOIP

We noted that HOIL1 deficiency resulted in a significantly greater defect in MDA5-driven IFN induction than did HOIP deficiency. We reasoned that the residual HOIL1 protein expressed in *Hoip*^-/-^ cells might be able to function independently of HOIP to support MDA5 signaling. Disruption of the *Hoip* gene occurred upstream of the region encoding the UBA that is required for interaction with the HOIL1 UBL domain (23, 55, 56), suggesting that a potential N-terminal fragment of HOIP in these cells would not be able to interact with and support HOIL1. We further tested this notion by transducing *Hoil1*^C458S/C458S^ MEFs with FLAG-tagged HOIL1 lacking its UBL domain (HOIL1[ΔUBL]; **Fig. 4A, B**). In these cells, endogenous HOIL1[C458S] formed a complex with and stabilized the expression of HOIP and SHARPIN (**Fig. 1A**, **2E** and **4B**) but lacked catalytic activity, whereas exogenous HOIL1[ΔUBL] was unable to interact with HOIP (**Fig. 4B**). HOIL1[ΔUBL] interaction with SHARPIN was reduced, but not completely absent, likely due to interaction between their LUBAC tethering motifs (LTMs) (55). HOIL1[ΔUBL] restored *Ifnb1* mRNA induction upon TMEV RNA transfection, similar to HOIL[WT], confirming that a direct interaction between HOIL1 and HOIP is not required for HOIL1 function in the MDA5 signaling pathway (**Fig. 4C**).

**Figure 4:**
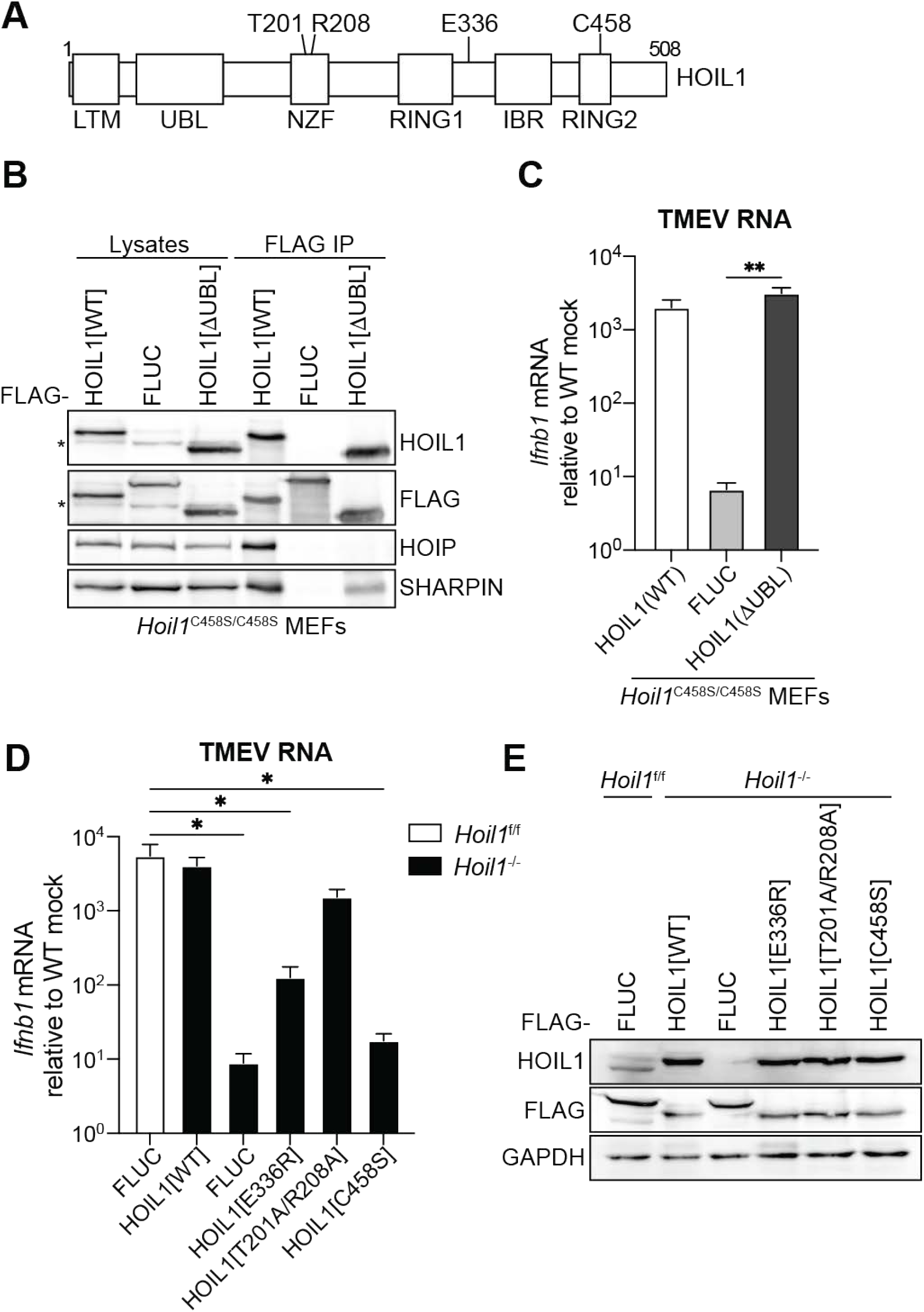
HOIL1 interaction with HOIP is dispensable for facilitation of MDA5 signaling. (**A**). Illustration of HOIL1 showing domain organization and key residues. LTM: LUBAC tethering motif; UBL: ubiquitin like domain; NZF: Npl4 zinc finger; RING; really interesting new gene; IBR: in-between RING domain. (**B-C**) *Hoil1*^C458S/C458S^ MEFs were transduced with 3xFLAG-HOIL1[WT], -HOIL1[ΔUBL], or –FLUC. (**B**) Immunoblot analysis of LUBAC subunit expression in whole cell lysates and after co-IP with the indicated FLAG-tagged proteins. (**C**) *Ifnb1* induction in the indicated MEF lines after TMEV RNA transfection. (**D-E**) *Hoil1*^-/-^ MEFs were transduced with 3xFLAG-tagged FLUC, - HOIL1[WT] or the indicated HOIL1 mutants. WT cells expressing 3xFLAG-FLUC were included as a control. (**D**) *Ifnb1* induction in the indicated MEF lines after TMEV RNA transfection. (**E**) Immunoblot analysis of expression of the indicated HOIL1 variants and FLUC in uninfected MEFs. For A and D, data shown are representative of 3 independent experiments. For B and C, data are combined from 3 independent experiments performed in duplicate, and the mean ± SEM is shown. Significance was determined by one-way ANOVA with Tukey’s multiple comparisons test. \**p*≤0.05, \*\**p*≤0.01.

These findings led us to investigate how HOIL1 may be recruited to targets of ubiquitination during MDA5 signaling, as LUBAC-independent roles of HOIL1 are not well described. We first assessed whether polyubiquitin binding by the NZF domain of HOIL1 is important for MDA5 signaling. However, disruption of HOIL1 NZF ubiquitin binding (HOIL1[T201A/R208A] (47, 57)) did not significantly impair IFN induction after TMEV RNA transfection (**Fig. 4D, E**). Two recent studies reported that the RING1, In-Between-RING (IBR) region and intervening linker region of HOIL1 also participate in the binding of K63- and M1-linked ubiquitin chains, and that this ubiquitin binding is important for allosteric activation of the HOIL1 catalytic site (44, 46, 47). We therefore examined whether polyubiquitin binding was important for MDA5 regulation by HOIL1 by mutating a key residue in HOIL1 (E336R, equivalent to E338R for human HOIL1) at the RING1-IBR linker-ubiquitin interface (47). Expression of HOIL1[E336R] in *Hoil1*^-/-^ MEFs only partially restored *Ifnb1* mRNA induction upon TMEV RNA stimulation (**Fig. 4D, E**), suggesting that binding of pre-existing polyubiquitin chains by the RING1-IBR domains is important for HOIL1 to regulate MDA5.

### HOIL1 ubiquitin ligase activity is necessary for MDA5-mediated MAVS activation

As HOIL1 ligase activity does not regulate RIG-I signaling, and since both RIG-I and MDA5 signal through MAVS, we reasoned that HOIL1 E3 enzymatic activity is likely required for MDA5-mediated activation of MAVS. Indeed, MAVS aggregation, a critical step for MAVS activation and innate immune signal propagation, was impaired in *Hoil1*^-/-^ MEFs during infection with mutEMCV, but not SeV (**Fig. 5A**). Expression of HOIL1[WT] restored MAVS aggregation upon mutEMCV infection, but HOIL1[C458S] did not (**Fig. 5B**). As MAVS is tethered to the outer mitochondrial membrane, we performed mitochondrial fractionation to assess the impact of HOIL1 ligase activity in the redistribution of MDA5 to mitochondria following stimulation by TMEV RNA. Endogenous MDA5 was enriched in the mitochondrial fraction upon viral RNA stimulation in cells expressing HOIL1[WT], but not in cells expressing HOIL1[C458A/S] (**Fig. 5C, D**). Collectively, these results indicate that HOIL1 E3 ligase activity promotes MDA5 translocation to the mitochondria and formation of the active MAVS signaling platform.

**Figure 5:**
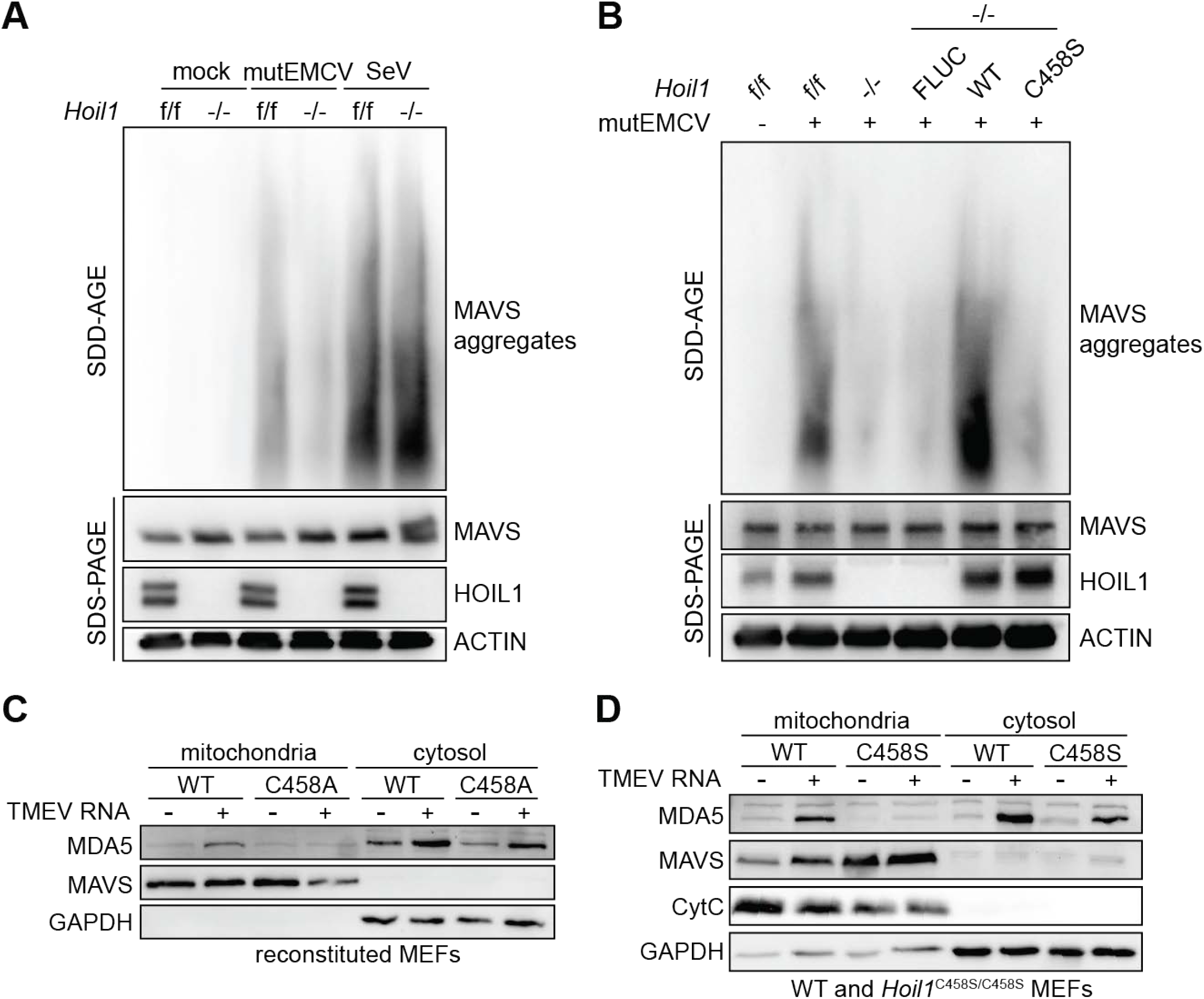
HOIL1 ubiquitin ligase activity promotes MDA5 interaction with and activation of MAVS. (**A**) WT and *Hoil1*^-/-^ MEFs were infected with mutEMCV (MOI of 0.5) or SeV (10 HAU/ml), or mock infected for 12 h. Cell lysates were subject to SDD-AGE (top) and SDS-PAGE (bottom) followed by western blot analyses to examine MAVS aggregation and total protein expression. (**B**) *Hoil1*^f/f^ (f/f) or *Hoil1*^-/-^ (-/-) MEFs complemented with FLUC, HOIL1[WT], or HOIL1[C458S] were infected with mutEMCV (MOI of 0.5, 12 hpi) and MAVS aggregation was examined as in A. (**C-D**) Immunoblot analysis of mitochondrial and cytosolic fractions from WT and *Hoil1*^-/-^ +HOIL1[C458A] MEFs (C), and WT and *Hoil1*^C458S/C458S^ MEFs (D) mock transfected or transfected with TMEV RNA (8 h) to examine MDA5 presence in the mitochondrial fractions after activation. Data shown are representative of at least 2 independent experiments.

### HOIL1 promotes ubiquitination of LGP2

Our data thus far indicate that HOIL1’s E3 ligase activity regulates an early, fundamental aspect of MDA5 activation. We posited that HOIL1 may modify MDA5 itself or, alternately, a key regulator that promotes MDA5 activation. Such regulators could modulate critical post-translational modifications of MDA5 (i.e. MDA5 ISGylation and K63-linked ubiquitination (58, 59)) or MDA5’s ability to bind dsRNA. Experiments using siRNA-mediated knock-down (KD) of endogenous *HOIL1* indicated that HOIL1 does not impact MDA5 ubiquitination or ISGylation in response to poly(I:C) stimulation (**Fig. 6A, B**). Given the crucial role of LGP2 in promoting MDA5 signaling (9, 12–14), we tested whether HOIL1 regulates MDA5 via LGP2. Co-immunoprecipitation of exogenously expressed proteins demonstrated that human HOIL1 could interact with both MDA5 and LGP2, but not with RIG-I (**Fig. 6C**). Ectopic expression of HOIL1 enhanced the ubiquitination of LGP2 (**Fig. 6D, E**) but not that of MDA5 or RIG-I (**Fig. 6D**). Conversely, KD of endogenous *HOIL1* reduced the ubiquitination of FLAG-tagged human LGP2 (**Fig. 6F**). Consistent with these data for ectopically expressed human LGP2, mutEMCV infection induced ubiquitination of endogenous LGP2 in MEFs in a HOIL1-dependent manner (**Fig. 6G**). Furthermore, while reconstitution of HOIL1-deficient MEFs with HOIL1[WT] fully restored LGP2 ubiquitination, HOIL1[C458S]-reconstituted cells showed only partially restored LGP2 ubiquitination during EMCV infection (**Fig. 6H**), suggesting that HOIL1 facilitates LGP2 ubiquitination through E3 ligase-dependent and -independent mechanisms. Notably, we were unable to detect linear polyubiquitination (Ub-M1) of LGP2, suggesting that HOIL1 is not acting through modulation of HOIP to facilitate ubiquitination of LGP2 (**Fig. 6E-G**). Together, these data support a model where HOIL1 regulates MDA5 activation through ubiquitination of LGP2.

**Figure 6:**
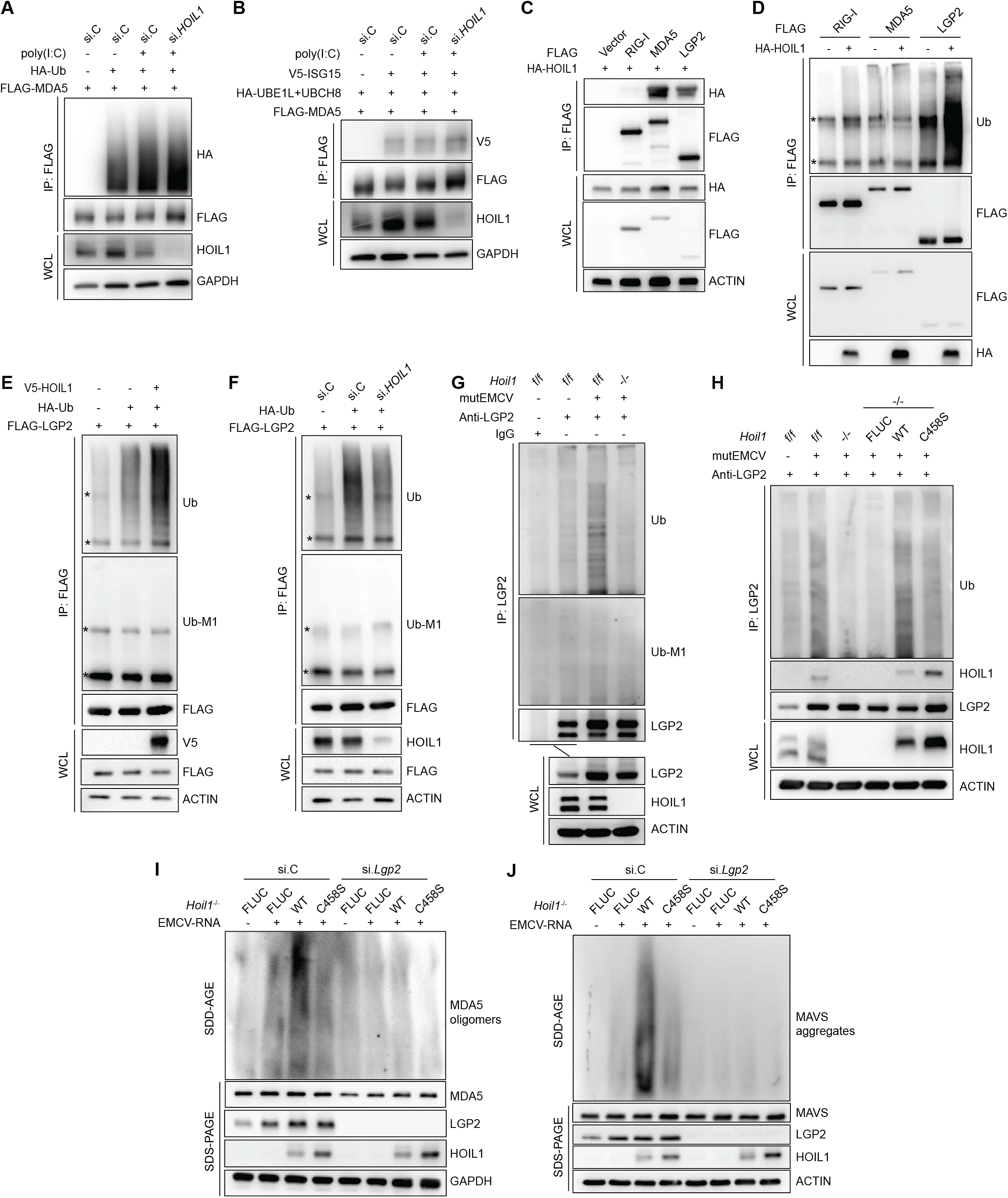
HOIL1 promotes ubiquitination of LGP2. (**A-B**) Ubiquitination (HA-Ub, A) or ISGylation (V5-ISG15, B) of FLAG-hMDA5 immunoprecipitated from 293T cells after siRNA-mediated knock-down (KD) of *HOIL1* (si. *HOIL1*) or control (si.C) and stimulation with poly(I:C) (1 ug/ml) for 6 h, or unstimulated. Expression of HOIL1 is indicated in the whole cell lysates (WCL). (**C**) Co-immunoprecipitation of HA-HOIL1 with FLAG-tagged RIG-I, MDA5 or LGP2 (IP: FLAG). Protein expression in WCL is shown below. (**D**) Ubiquitination of FLAG-tagged RIG-I, MDA5 or LGP2 IP’d from 293T cells with or without overexpression of HA-HOIL1 (top). Amounts of FLAG-tagged proteins IP’d or in WCL are shown below. (**E-F**) Total ubiquitination (top) and linear ubiquitination (Ub-M1, middle) of LGP2 in the presence or absence of exogenous HA-ubiquitin and V5-HOIL1 (E) or after KD of *HOIL1* (F) in 293T cells. (**G**) Ubiquitination of endogenous LGP2 IP’d from WT (f/f) and *Hoil1*^-/-^ MEFs with and without mutEMCV infection (MOI of 0.1, 12 hpi). Total ubiquitination (top) and linear ubiquitination (middle) are shown. (**H**) Ubiquitination of endogenous LGP2 IP’d from WT (f/f) and *Hoil1*^-/-^ MEFs complemented with FLUC, HOIL1[WT] or HOIL1[C458S], with and without mutEMCV infection (MOI of 0.1, 12 hpi). (**I-J**) SDD-AGE western blot analyses of MDA5 oligomers (I) and MAVS aggregates (J) induced by EMCV-RNA (400 ng/ml for 16 h) in *Hoil1*^-/-^ MEFs complemented with FLUC, HOIL1[WT] or HOIL1[C458S], after control (si.C) or *Lgp2* (si.*Lgp2*) KD. Total protein expression is shown by SDS-PAGE western blot analyses below. Asterisks (*) in panels D-F indicate non-specific bands. Data shown are representative of at least 2 independent experiments.

To corroborate this proposed model, we silenced endogenous *Lgp2* in *Hoil1*^-/-^ MEFs reconstituted with HOIL1[WT] or HOIL1[C458S] (or FLUC [control]), and examined the formation of MDA5 oligomers and MAVS aggregates in response to EMCV RNA stimulation (**Fig. 6 I, J**). Knock-down of *Lgp2* almost completely abrogated the formation of MDA5 oligomers and MAVS aggregates in all cell lines, consistent with other studies demonstrating that LGP2 facilitates MDA5 oligomerization and thereby downstream signaling ((9, 13, 14)). MDA5 and MAVS higher-order assemblies were restored by HOIL1[WT] expression in control siRNA (si.C) transfected cells, but not in *Lgp2*-depleted cells, indicating that HOIL1 cannot facilitate MDA5 oligomerization in the absence of LGP2. As compared to cells expressing HOIL1[WT], HOIL1[C458S] expression only slightly promoted MDA5 and MAVS higher-order assembly formation in si.C-transfected cells. Further, HOIL1[C458S] expression did not induce MDA5 or MAVS aggregate formation in *Lgp2*-silenced cells (**Fig. 6I, J**). These results indicate that MDA5 oligomerization, and ensuing MAVS activation, are largely dependent on HOIL1’s ligase activity in a manner dependent on LGP2.

Altogether, our data show that HOIL1 promotes MDA5 activation and downstream signaling through mediating LGP2 ubiquitination. Our data further indicate that HOIL1-mediated ubiquitination of LGP2 occurs through HOIL1 ligase-dependent and - independent mechanisms, unveiling a novel role for HOIL1 in antiviral immunity.

## Discussion

MDA5 antiviral signaling initiates the innate immune response to a range of RNA viruses and for positive outcomes of viral disease. Beyond antiviral immunity, aberrant MDA5 activation has been identified as a driver of inflammation in patients with Singleton-Merten syndrome and Aicardi-Goutières syndrome, and implicated in the pathogenesis of other immune disorders like type 1 diabetes, systemic lupus erythematous, and ADAR1 deficiency (16–19, 60–62). Thus, an in-depth understanding of the mechanisms controlling MDA5 signaling is important for developing more targeted immunomodulatory strategies to enhance antiviral immunity or limit auto-inflammation.

In this study, we resolved the role of HOIL1 in MDA5 antiviral signaling by delineating the relative contributions of HOIL1 E3 ubiquitin ligase activity and LUBAC activity. The role of the LUBAC in RIG-I signaling has been previously studied, with conflicting conclusions likely stemming from differences in HOIP expression levels, incomplete deletion of LUBAC components, and cell type-dependent effects (20, 33–37). Nevertheless, multiple groups have reported a partial defect in RIG-I-mediated IFN induction with loss of HOIP, or HOIP’s E3 catalytic activity (36, 37), similar to what we present in this study. We extend these findings by identifying a similar role for HOIP ligase activity in MDA5 signaling. These findings suggest that the LUBAC regulates a component(s) common to MDA5 and RIG-I signaling, likely at the level of MAVS, or downstream of MAVS, where the signaling by both RLRs converges. This is in line with findings by Liu *et al*. (36) and Teague *et al*. (37), which support a model in which linear ubiquitin chains generated by HOIP facilitate the recruitment and retainment of NEMO to the MAVS signaling complex, which in turn recruits and activates kinases that activate IRF3 and NF-κB. Liu *et al*. also reported possible redundancy between the LUBAC and TRAF2 in promoting MAVS signaling, which may partly explain why the LUBAC is not absolutely required for RLR-mediated IFN induction. Nevertheless, given the multiple proposed cellular functions of LUBAC, additional studies are warranted to further define the mechanisms by which LUBAC regulates RLR signaling, and their respective contributions of these mechanisms (upstream and downstream of MAVS) to the effects seen on RLR signaling.

Recently, HOIL1 has been shown to be able to catalyze ester-linked ubiquitination of serine and threonine residues on substrates and within ubiquitin itself, but the range of substrates and the physiological functions of ester-linked ubiquitination identified to date are limited (40–45, 52, 63, 64). Importantly, here we demonstrated that HOIL1 ubiquitin ligase activity is critical for MDA5 antiviral signaling after stimulation with multiple viruses and viral RNAs, including SARS-CoV-2, and that HOIL1 can regulate MDA5 independent of its association with the LUBAC. A recent study indicated that HOIL1 and SHAPRIN can form homodimers and heterodimers without HOIP (65), and other LUBAC-independent roles have been reported (66). However, the polyubiquitin binding region between the RING1 and IBR domains of HOIL1 was shown to be important for IFN induction, indicating that the generation of M1-polyubiquitin by HOIP, or K63-polyubiquitin by another E3 ligase is likely important for HOIL1 function.

Our data indicate that HOIL1 acts upstream of MAVS, at the level of MDA5, and enhances MDA5 oligomerization during infection. Further, we have identified LGP2, an important positive regulator of MDA5 signaling (9, 12, 13), as a target of ubiquitination by HOIL1. However, HOIL1 appeared to promote ubiquitination of LGP2 directly through its own ligase activity as well as indirectly, perhaps by facilitating recruitment of another E3 ligase to LGP2. Although HOIL1 interacts with HOIP, we were unable to detect changes in linear ubiquitination of LGP2, suggesting that the ubiquitination catalyzed on LGP2 is independent of HOIP. Although the exact mechanism(s) of how LGP2 potentiates MDA5 signaling is still elusive, it has been shown that LGP2 promotes ATP-dependent nucleation of MDA5 filament assembly from internal RNA sites (9, 67). LGP2 exhibits higher affinity for dsRNA than MDA5 (9, 68, 69), suggesting that LGP2 binds viral RNAs before recognition by MDA5. Future studies will be needed to determine the manner in which LGP2 is ubiquitinated, whether another E3 ligase is involved, and how LGP2 ubiquitination impacts MDA5 oligomerization and thereby signaling at the molecular level. Overall, our study identifies a novel function for the HOIL1 E3 ubiquitin ligase as critical regulator of MDA5 activation and IFN induction.

## Acknowledgements

We thank: B. Studnicka, M. Byrne, J. Marshall, F. Faraji, and the UIC Research Resources Center Genome Core for technical support; S. Whelan and N. Sarute for VSV-GFP; H.W. Virgin for MNoV CR6; H. Lipton for TMEV GDVII; F. J. M. van Kuppeveld for mutEMCV; J. Schoggins and C. Rice for lentiviral vectors; J. U. Jung for HA-hHOIL1 and V5-hHOIL1 expression vectors; P. Cohen for the *Hoil1*^C458/C458^ KI mice; Center d’ImmunoPhenomique (Ciphe) for providing the mutant mouse line (Allele: Rbck1^tm1a(EUCOMM)Hmgu^), INFRAFRONTIER/EMMA (www.infrafrontier.eu, PMID: 25414328), and Institut de Transgenose (INTRAGENE, Orleans, France) from which the mouse line was distributed (EM:09852) (70, 71). Associated primary phenotypic information may be found at www.mousephenotype.org.

## Funding

This work was funded by NIH/NIAID R01 AI150640 and by UIC institutional start-up funds to D.A.M. This work was also supported in part by NIH/NIAID R37 AI087846 (to M.U.G.).

## Materials and Methods

### Plasmids and cloning

*Hoip* (*Rnf31*) cDNA was PCR amplified from mouse cDNA, and FLUC (firefly luciferase) was amplified from pTrip-CMV-FLUC-EYFP (72) and cloned into p3xFLAG-CMV-10 (Sigma) NotI/XbaI. 3xFLAG-*Hoil1* (*Rbck1*) cDNA was synthesized as a gene block (IDT). FLAG-tagged constructs were PCR amplified with Gateway adapters and cloned into pDONR221 using BP Clonase (Invitrogen). *Hoil1* and *Hoip* mutations and deletions were generated using the Q5 site-directed mutagenesis kit (New England Biolabs). FLAG-tagged cDNAs were transferred into pTrip-CMV-Puro-T2A-RFP (72) (HOIL1 and FLUC) or pSCRBBL (a derivative of pSCRPSY carrying the blasticidin resistance gene; HOIP and FLUC) lentiviral vectors using LR Clonase II (Invitrogen). LentiCRISPR v2 (a gift from Feng Zhang (Addgene plasmid # 52961) (73)) expressing gRNA complementary to *Hoip* exon 3 was created by synthesizing complementary DNA oligos (IDT) with the sequence GATGGATTGAGTTTCCCCGA and strand-specific overhangs compatible with ligation after vector digestion with BsmBI. Oligonucleotides were annealed and ligated with lentiCRISPR v2 plasmid digested with BsmBI. Complementation of *Hoip*^-/-^ cells was achieved by introducing synomymous mutations across the gRNA recognition site in *Hoip* cDNA. FLAG-hMDA5, FLAG-hRIG-I, HA-Ub, pCAGGS–HA–Ube1L, pFLAG–CMV2–UbcH8 and V5-hISG15 have been described previously (59, 74). Human LGP2 was amplified from human cDNA and cloned into pcDNA3.1-FLAG. HA-hHOIL1 and V5-hHOIL1 were kindly provided by J. U. Jung (Cleveland Clinic) and have been described previously (34).

### Viral stocks

Stocks of MNoV strain CR6 were generated from a molecular clone as previously described (75, 76). Briefly, a plasmid encoding the CR6 genome was transfected into 293T cells to generate infectious virus, which was subsequently passaged on BV2 cells. After two passages, BV2 cultures were frozen and thawed to liberate virions. Cultures were cleared of cellular debris, then concentrated by filtration though a Vivaflow 200 PES filter with a 10,000 molecular weight cut-off (Sartorius). Titers of MNoV stocks were determined by plaque assay on BV2 cells (75, 76). Sendai virus Cantell strain was obtained from ATCC (VR-907) and used to infect BMDCs directly. VSV-GFP (77) and TMEV strain GDVII were propagated in Vero cells and BHK cells, respectively, and purified by filtration of cell culture supernatant through a 0.22 μm filter. Titers of VSV-GFP and TMEV stocks were determined by plaque assay on Vero cells and BHK cells, respectively. mutEMCV, which carries two point mutations (C19A/C22A) in the zinc domain of the L protein, was kindly provided by F. J. M. van Kuppeveld (Utrecht University) and was propagated in BHK-21 cells (50). To isolate viral RNA replication intermediates, BHK cells were infected with TMEV or VSV-GFP at an MOI of 0.05 and incubated at 37°C with 5% CO_2_. At 21 hpi, infected cells were lysed with TRI-Reagent and RNA isolated according to the manufacturer’s instructions. EMCV-RNA, SARS-CoV-2 RNA, and RABV_Le_ RNA were generated as previously described (59, 74, 78).

### Cell culture, treatments and infections

Mouse embryonic fibroblasts (MEFs) and HEK-293T cells were cultured in DMEM (Corning) supplemented with 10% FBS, 1% L-glutamine, 0.5% penicillin/streptomycin at 37°C with 5% CO_2_. MEFs were isolated from E14 embryos and transformed with SV40 large T-antigen. Immortalized *Hoil1*^f/f^ MEFs were transiently transfected with a plasmid expressing Cre-recombinase (pTrip-CMV-Cre-Puro-T2A-RFP) using *Trans*IT-LT1 (Mirus), and *Hoil1*^-/-^ and *Hoil1*^f/f^ single-cell clones isolated and identified by PCR. To generate *Hoip*^-/-^ MEFs, WT MEFs were transduced with LentiCRISPRv2 expressing gRNA targeting *Hoip* exon 3 and selected with puromycin. Single cell clones were evaluated by western blot and sequencing to identify knockout populations.

For lentiviral transductions, HEK293T cells were transfected with lentiviral expression plasmids, along with pMD2.D and psPAX2 lentiviral packaging plasmids, using *Trans*IT-LT1 (Mirus). Supernatants were collected at 24 and 48 h, filtered through a 0.22 μm filter and added to immortalized MEFs. The next day, transduced cells were selected with puromycin or blasticidin until all untransduced control cells were dead.

Transient gene knockdown in HEK293T and MEFs was performed using SMARTpool small interfering (si)RNAs (Horizon Discovery) as previously described (79). These include specific siRNAs targeting human *HOIL1* (L-006932-00-0005), mouse *Lgp2* (L-050413-00-0005), and Non-targeting Control Pool (D-001810-10-05).

For viral infections, immortalized MEFs were plated in tissue-culture treated plates, and 20 h later infected with SeV (MOI of 3), VSV-GFP (MOI of 3), TMEV GDVII (MOI of 3), or mutEMCV (MOI of 0.5) for the indicated times. Alternatively, viral RNA isolated from infected BHK cells was transfected into cells at a concentration of 500 µg/ml using *Trans*IT-mRNA transfection kit (Mirus). RABV_Le_ RNA (2 pmol/ml) was transfected with Lipofectamine RNAiMAX Transfection Reagent (Invitrogen) as previously described (59).

To assess sensitivity to TNFα-induced cell death, immortalized MEFs were plated in tissue-culture treated plates and 20 h later treated with 150 μg/ml TNFα. At 48 h, the medium was removed, and the cells fixed and stained with 0.1% crystal violet in 20% ethanol for 15 min, then gently washed with water and imaged.

Bone marrow-derived dendritic cells (BMDCs) were differentiated in RPMI-1640 (Corning) supplemented with 10% FBS, 1% L-glutamine, and 2% conditioned media from J558L cells containing GM-CSF in a 37°C, 5% CO_2_, humidified incubator for 7 days. For RNA and protein analyses, non-adherent cells were harvested and infected with MNoV CR6, TMEV GDVII or VSV-GFP at a multiplicity of infection (MOI) of 3, with SeV at an MOI of 0.3 (0.3 EID_50_/cell), mock infected, or treated with 50 ng/ml LPS (E. coli O55:B5, MilliporeSigma), and re-plated in tissue culture-treated dishes.

At the indicated time points, cells were lysed in TRI-Reagent (MilliporeSigma) and stored at -80°C for RNA extraction, or in 2x Laemmli buffer, boiled for 10 minutes and stored at -20°C for immunoblot analyses.

### Mice

B6NCrl;B6N-A^tm1Brd^Rbck1^tm1a(EUCOMM)Hmgu^ mice were acquired from INFRAFRONTIER/EMMA (European mutant mouse archive) (70, 71) and have been described previously (48). *Mda5*^-/-^ (B6.Cg-Ifih1tm1.1Cln/J, stock #015812) mice were purchased from Jackson Laboratories (Bar Harbor, ME). *Hoil1*^C458S/C458S^ KI mice were a gift from Philip Cohen (41) and were generated by Taconic Artemis. Mice were housed at University of Illinois Chicago under specific pathogen-free conditions in accordance with Federal and University guidelines, and protocols were approved by the University of Illinois Chicago Animal Care Committee.

### RNA isolation and quantitative reverse transcription-PCR

RNA was isolated according to the manufacturer’s instructions. RNA samples were treated with Turbo DNA-free DNase (Invitrogen) and 1 μg of RNA used as a template for cDNA synthesis with random primers and ImProm-II reverse transcriptase (Promega). Quantitative PCR (qPCR) was performed on a QuantStudio 3 Real-Time PCR System (Applied Biosystems) using Amplitaq Gold (Applied Biosystems) and predesigned probe-based assays for: *Ifnb1* (Mm.PT.58.30132453.g), *Ifit2* (Mm.PT.58.28800045.g), *Il6* (Mm.PT.58.10005566) (IDT). *Rps29* and MNoV TaqMan qPCR assays were performed as described previously (20, 80). MNoV genome copies and mRNA levels were determined relative to standard curves and normalized to *Rps29* using standard curves or by ΛΛC_T_ relative to mock.

### Co-immunoprecipitations

For IP of the LUBAC complex, MEFs were lysed in 1% NP-40, 150 mM NaCl, 50 mM Tris-HCl (pH 7.5), 1 mM EDTA. Lysates were centrifuged at 10,000g for 12 min and supernatant incubated with anti-FLAG magnetic agarose and further processed according to manufacturer’s instructions (Pierce A36797). For endogenous LGP2 IP, MEFs were infected with mutEMCV (MOI of 0.1) for 12 hours, and cell lysates were prepared and precleared with Protein G Dynabeads (Invitrogen) at 4°C for 2 hours. The lysates were then incubated with Protein G Dynabeads pre-conjugated with anti-LGP2 (Proteintech, 11355-1-AP) or normal rabbit IgG (CST, 2729S) at 4°C for 4 hours. The beads were washed four times with RIPA buffer (20 mM Tris-HCl, pH 8.0, 150 mM NaCl, 1% (v/v) NP-40, 1% (w/v) deoxycholic acid, and 0.01% (w/v) SDS), and proteins were eluted in 1× Laemmli SDS sample buffer. The protein samples were separated on Bis–Tris SDS-PAGE gels, transferred onto PVDF membranes (Bio-Rad), and analyzed by immunoblotting.

### Mitochondrial fractionation

Twenty million cells were trypsinized, pelleted, and washed three times with PBS. Cell pellets were resuspended in 1 ml Buffer A (10 mM Tris-HCl pH 7.5, 10 mM NaCl, 1.5 mM MgCl_2_, protease inhibitors) for 10 min on ice. Cell suspensions were passed through a 26 G needle 10 times, transferred to a dounce homogenizer, and homogenized for 30 strokes. 600 µl of Buffer B (525 mM mannitol, 175 mM sucrose, 12.5 mM Tris-HCl pH 7.5, 2.5 mM EDTA) was then added to the lysates, and the mixture was centrifuged at 1500 g for 5 min at 4°C to pellet debris. Centrifugation was repeated until there was no visible pellet. The supernatant was then centrifuged at 7500 g for 15 min at 4°C to pellet mitochondria and mitochondrial-associated membranes. Pellets were resuspended in 1 ml Buffer C (10 mM Tris-HCl pH 7.5, 0.15 mM MgCl_2_, 0.25 M sucrose) and centrifuged at 9500 g for 10 min. The wash with Buffer C was repeated and pellets were lysed with a 1:1 mixture of RIPA buffer and 2x Laemmli Buffer and heated at 98°C for 5 min. This protocol was adapted from (81).

### Semi-denaturating detergent agarose gel electrophoresis (SDD-AGE)

MDA5 oligomerization and MAVS aggregation in MEFs that were transfected with RNA ligands or infected with viruses was assessed by SDD–AGE as previously described (59, 82). Briefly, to assess MAVS aggregation, MEFs were infected with mutEMCV (MOI of 0.5) or SeV (10 HAU/ml) for 12 h, or transfected with EMCV-RNA (400 ng/ml) for 16 h. After treatment, cells were harvested and suspended in buffer A (10 mM Tris-HCl, pH 7.5, 1.5 mM MgCl_2_, 10 mM KCl, 0.25 M D-mannitol), followed by cell lysis through grinding. Cell lysates were then centrifuged at 700 ×*g* for 10 min to remove cell debris. The supernatants were transferred to a new tube and further centrifuged at 10,000 ×*g* for 30 min at 4°C to separate the crude mitochondria from cytosolic extracts. The crude mitochondria were resuspended in 1× sample buffer (0.5× TBE, 10% glycerol, 2% SDS, and 0.0025% bromophenol blue) and loaded onto a 1.5% agarose gel. Electrophoresis was performed at a constant voltage of 100 V at 4°C until the bromophenol blue dye reached the bottom of the agarose gel. Proteins were then transferred onto a PVDF membrane and analyzed by immunoblotting using the specified antibodies.

### IRF3 dimerization

Endogenous IRF3 dimerization was determined by native PAGE as previously described (83). Briefly, MEFs (in a 6-well plate) that were infected with mutEMCV (MOI of 5) or transfected with RABV_Le_ RNA (5 pmol/ml) for 8 h were scraped off in ice-cold PBS, pelleted, and lysed in 60 μl NP-40 buffer containing 10% glycerol. Cell lysates were then subject to three freeze-thaw-vortex cycles, followed by centrifugation at 16,000 ×*g* at 4°C for 20 min. The cleared lysates were mixed with 2× native-PAGE loading buffer and separated on a 7.5% acrylamide/bis-acrylamide (37.5:1) Tris-Glycine gel with 1% sodium deoxycholate added to the inner chamber buffer. Proteins were then transferred onto a PVDF membrane and analyzed by immunoblotting.

### Immunoblot analyses

Proteins were separated by SDS-PAGE or SDD-AGE and transferred to PVDF membrane, blocked with 5% non-fat dry milk in TBS containing 0.1% Tween 20. Membranes were incubated with the primary antibodies against: HOIL1 (Santa Cruz Biotechnologies, sc393754), HOIP (Abcam, ab46322), SHARPIN (Proteintech, 14626-I-AP), FLAG (M2, Sigma), MAVS (Cell Signaling Technologies (CST), 4983S), MDA5 (Adipogen, AL180), LGP2 (Proteintech, 11355-1-AP), IRF3 (CST, 11904S), IRF3-pS396 (CST, 29047S), cytochrome c (CST, 11940S), HA (CST, 3724S), V5 (CST, 13202S), ubiquitin (Santa Cruz Biotechnologies, sc-8017), M1-ubiquitin (Sigma-Aldrich, MABS451), β-actin (Sigma), GAPDH (CST, 97166S), followed by incubation with goat anti-rabbit or anti-mouse IgG-HRP. Blots were incubated with Immobilon Western Chemiluminescent HRP Substrate (Millipore) and imaged on a ChemiDoc Imaging System (Bio-Rad).

### Statistical analyses

Data were analyzed with Prism 10 software (GraphPad Software, San Diego, CA) as described in the figure legends.

## Notes

### Competing Interest Statement

The authors have declared no competing interest.

